# A network regularized linear model to infer spatial expression pattern for single cell

**DOI:** 10.1101/2024.11.20.624541

**Authors:** Chaohao Gu, Hu Chen, Zhandong Liu

## Abstract

Spatial transcriptomics, situated at the intersection of genomics and spatial biology, offers profound insights into the spatial organization of gene expression within tissues. However, its potential has been constrained by either limited resolution or throughput. While single-cell RNA-seq allows for in-depth profiling of cellular gene expression, the crucial spatial information is often sacrificed during sample collection. In a groundbreaking fusion of these two techniques, our research introduces the glmSMA computational algorithm. This innovative approach aims to predict cell locations by integrating scRNA-seq data with spatial-omics reference atlases. The essence of glmSMA lies in formulating cell mapping as a convex problem, strategically minimizing differences between cellular expression profiles and location expression profiles through L1 and Generalized L2 regularization. Our algorithm has undergone rigorous testing across diverse tissues, including mouse brain, drosophila embryo, and human PDAC samples. The compelling results validate glmSMA’s efficacy, demonstrating its capability to faithfully recapitulate spatial gene expression and anatomical structures. This marks a significant stride forward in overcoming the limitations of existing spatial transcriptomic techniques.

## Background

Precise spatial regulation of gene expression at the individual cell level plays a crucial role in normal development and disease pathogenesis. A notable example is the spatial expression of the Hippo signaling pathway during early embryogenesis in Drosophila, which is essential for orchestrating the correct patterning of asynchronous cell proliferation^1,2^. While single-cell RNA sequencing (scRNA-seq) has revolutionized the measurement of a vast number of genes in individual cells, the inherent challenge lies in the loss of spatial information post-tissue digestion and cell dispersion^3^.

Spatial-omics techniques today fall into two main categories: Next-Gen Sequencing (NGS)-based methods and imaging-based methods. Examples of NGS-based methods include Slide-seq and spatial transcriptomics. Slide-seq offers genome-wide measurements for obtaining spatially resolved gene expression data at the single-cell level^4^. However, it comes with limitations, such as a low gene capture rate and relatively small barcoded bead size, resulting in a confined measurable area. On the other hand, spatial transcriptomics, exemplified by technologies like 10x Visium, provides high-quality mRNA transcripts over a larger area but at the cost of lower resolution compared to Slide-seq^5^. In addition, image-based methods resemble the fluorescence in-situ hybridization (FISH) technique^6^. FISH-like methods offer high-resolution mRNA level information in a tissue of interest. However, they often face constraints in the number of genes that can be measured. Navigating the strengths and limitations of these spatial-omics techniques is pivotal for advancing our understanding of spatial gene expression regulation in diverse biological contexts.

The seamless integration of single-cell and spatial transcriptomics (ST) data holds the promise of characterizing the spatial distribution of the entire transcriptome at single-cell resolution, harnessing the complementary insights offered by each. However, current integration methods fall short of providing satisfactory outcomes in real data analysis. Several cell-type deconvolution methods for ST data, including RCTD^7^, Cell2location^8^, and MIA^9^, exist; however, when applied to sequencing-based ST data, they merely estimate cell type proportions within spatial spots, lacking the capability to achieve true single-cell resolution. In addition to established NGS-based methods like Slide-seq and spatial transcriptomics, alternative approaches such as novosparc^10^, DistMap^11^, and CellTrek^12^ contribute to the landscape of spatial-omics techniques. However, each method exhibits its own set of limitations. CellTrek^12^, for instance, faces a challenge in achieving high recall due to its automatic removal of at least 30% of cells with low confidence scores. This trade-off between precision and recall can impact the comprehensive capture of spatial information. Novosparc^10^, while promising, has primarily undergone testing in simulated datasets, raising questions about its real-world applicability and performance on authentic biological samples. DistMap^11^, while effective, is constrained by its suitability for simple tissues. This limitation stems from the oversimplification of gene expression through binarization, which may compromise its efficacy in capturing the complexity of spatial regulation in more intricate biological systems. Consequently, the persisting limitation of not attaining single-cell resolution remains unaddressed. In the context of image-based ST data, methods designed to infer unmeasured gene expressions, such as Tangram^13^, SpatialScope^14^, and CARD^15^, exhibit insufficient accuracy, particularly in the presence of sparse ST expression data. Thus, there persists a crucial need for precise statistical and computational methods to effectively bridge the gap between single-cell and ST datasets.

## Methods

GlmSMA represents a groundbreaking integration of spatial transcriptomics and single-cell RNA sequencing (scRNA-seq) to predict cell locations within a spatial reference. This innovative approach involves constructing a linear regression model incorporating L1 norm and generalized L2 norm. There is a linear correlation between expression distance and physical distance. Within a given anatomical structure, cells in closer proximity exhibit more similar expression patterns. The model aims to estimate the linear relationship between the location expression and cellular expression of specific marker genes, where the generalized L2 norm is induced by the graph Laplacian matrix calculated from spatial coordinates derived from the spatial reference. To achieve a balanced approach in selecting locations for single cells within limited areas, glmSMA leverages L1 and generalized L2 regularization. The accuracy of cell location prediction largely depends on the choice of marker genes. To identify these markers, we implemented three alternative strategies: (1) selecting highly variable genes, (2) choosing spatially informative genes based on Moran’s I scores, and (3) identifying anchor genes shared between the scRNA-seq and spatial transcriptomics datasets. Any of these methods can be used to select marker genes for prediction. Moreover, researchers have the flexibility to input their own marker gene list. In essence, glmSMA minimizes the location and cellular expression of marker genes, resulting in a focused selection of potential locations for each individual cell.

Validating the predicted cell locations poses a challenge due to the absence of ground truth in the experimental procedure. Direct ground truth for cell locations is often unavailable in scRNA-seq data due to experimental constraints. To address this, we adopted alternative strategies for estimating ground truth in each dataset. First, the Kullback-Leibler divergence (KL) is calculated between the original cell type distribution based on spatial transcriptomics and the predicted distribution. Lower KL scores indicate more accurate predictions. Second, Pearson’s correlation is computed on the spatial expression of marker genes between the reference and reconstructed patterns derived from cell mapping results. This analysis reveals a strong positive correlation in multiple tissues. Third, manual inspection of cell type distribution based on anatomical structures further contributes to the validation process. Through these comprehensive approaches, glmSMA offers a robust means of validating cell mapping results despite the absence of ground truth information.

### 2.1 Objective function to minimize

Suppose that the reference atlas has n positions with p genes, and the scRNA-seq data set has m cells with the same number of p genes (usually n > m). We aimed to assign the m cells into n positions using linear regression model with L1 norm and generalized L2 norm via graph Laplacian. The initial step is to create the position to gene expression matrix and the cell to the gene expression matrix. Based on the two matrices, we create two graphs – position to position graph and cell to cell graph. If the Euclidean distance between two locations is lower than a specific threshold and the two locations belong to a same anatomical structure, the two locations are connected in the first graph. If the similarity of gene expression levels between two cells is lower than a specific threshold, the two cells are connected in the cell-to-cell graph. Then we create two random walk normalized graph Laplacian matrices, which encourage smoothness on coefficients that are connected in the two graphs.

Our model uses a linear method to measure the differences in gene expression levels in assigning cells to locations. The optimal solution minimizes the differences between gene expression levels of individual cells and gene expression levels of locations. For each individual cell, we want to minimize the following objective function,

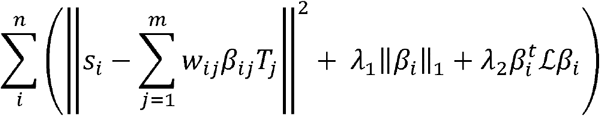

Where *s* ∈ *R*^*n*x*g*^ is the single-cell expression matrix (n is the number of cells, g is the number of genes),*T* ∈ *R*^*m*x*g*^ is marker genes expression matrix of reference atlas (g is number of genes, and m is the number of locations in the reference atlas), ℒ is normalized graph Laplacian matrix computed from the location distance graph, and *w* ∈ *W*^*i* x*j*^ is the personalized weight matrix for each location in each cell. The L1 norm penalization encourages sparsity on the coefficients, which guarantees that one cell can only be assigned into a small number of locations. The generalized L2 norm encourages the smoothness of the coefficients, which guarantees that cells with similar gene expression leve ls are more likely to be assigned into closer locations and one cell assignment is more likely to form a patch instead of scattered distribution. Overall, our method makes full use of the relationship between cell-to-cell physical distance and cell to location expression distance.

### 2.2 Data analysis

For the Slide-seq data, we normalized the data in log-space following the previous pipeline [6]. Let *d*_*ij*_ be the raw count for gene i in cell j; we normalized it as

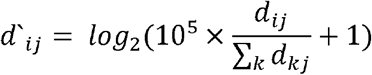

Then we down-sampled the Slide-seq datasets by rounding the physical location coordinates to the next integer multiple of 50 and filtered out low quality locations, where positions with less than 50 genes expressed were discarded, resulting in 7,724 cells in the mouse cerebellum section and 7,790 cells in the mouse hippocampus section. Then we used summed transcriptomes of neighboring locations as the new expression profiles in the rounded locations.

To find the highly variable genes in the Slide-seq dataset, Raw counts in each cell were first normalized by the total counts and scaled by the median number of raw counts across all cells. For each cell we added a pseudocount of 1 before log transformation. Highly variable genes were selected based on the Fano factor, defined as 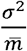, where σ^2^ is the variance of gene expression and 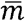 is the mean of expression across all cells. A certain mean threshold will be chosen for the Fano factor. For the 10x Visium spatial transcriptomics data, we use the Seurat package to find the highly variable genes.

Next, we built a graph Laplacian matrix based on the spatial coordinates and anatomical structure. In physical space, we first constructed an undirected graph G = (V, E), where V is a set of nodes consisted of locations in the reference atlas and E is a set of edges consisted of the Euclidean distance between locations. Two locations are connected if the Euclidean distance between them is less than a selected threshold and both locations are in same anatomical structure. The random-walk normalized Laplacian matrix on physical locations is defined as,

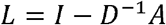

where D is the degree matrix and A is the adjacency matrix in graph G.

The elements of L are shown as,

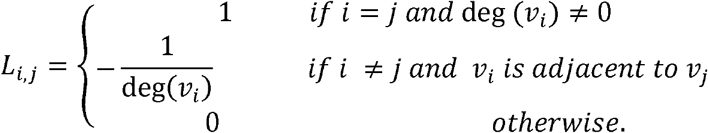

### 2.3 Selection of regularization penalties

L1 norm encouraged the sparsity of the cell mapping and generalized L2 norm encouraged the smoothness of the cell mapping. If we want to map the cells to a large domain, we can increase the L2 parameters and decrease the L1 parameters. Also, we can increase L1 parameter to map cells to several sparse locations. For the simulated dataset, we used a stepwise increasement for the L1 and L2 parameters and found the optimal solution according to the highest prediction accuracy for the cell locations.

### 2.4 glmSMA pipeline

To reconstruct spatial expression patterns, our algorithm performs the following steps:

1. Read the gene expression matrix from scRNA-seq and location matrix from reference atlas
2. Optional:
  a. Find the spatial differential expressed genes based on the Moran’s I scores.
  b. Find the highly variable genes
  c. Find anchor genes from spatial reference atlas based on the scRNA-seq data
  Then the input will be altered based on the genes of interest.
3. Construct the normalized graph Laplacian matrix based on the anatomical structures and Euclidean distance between locations.
4. Optional: Set personalized weight matrix for each location and cell.
5. Using glm to solve the convex function with L1 norm and generalized L2 norm.
6. Assign cells to targeting locations based on the distribution of *β* in the objective function. For each cell, output a probability vector across all spatial locations.
7. Reconstruct the spatial patterns based on the expression profiles in the scRNA-seq data and cell locations from the mapping results.

## Results

### Overview of glmSMA

Presenting glmSMA, a novel linear regression model designed to integrate single-cell RNA sequencing (scRNA-seq) reference data with spatial transcriptomics (ST) data across various experimental platforms (Fig.1; Methods). glmSMA seamlessly accommodates both sequencing-based ST data (such as 10x Visium and Slide-seq) and image-based ST data, making it a versatile tool for spatial analysis. The model combines L1 and generalized L2 norm regularization, enabling it to resolve spot-level data containing multiple cells down to single-cell resolution—particularly excelling with sequencing-based ST data. The L1 norm induces sparsity in single-cell distribution, ensuring an efficient and focused representation of individual cells. Meanwhile, the generalized L2 norm, informed by a graph Laplacian constructed from anatomical structures and spatial coordinates, promotes smoothness in cell distribution within local regions16. This dual regularization approach allows glmSMA to accurately assign single cells to a limited number of small regions at single-cell resolution, offering a refined understanding of spatial organization. For each cell, glmSMA generates a probability vector across all spatial locations. By default, we report the top five most probable locations for each cell. Through rigorous benchmarking on both simulated and in situ datasets, glmSMA has demonstrated robust performance, accurately mapping cells across diverse tissue types. Its application to various samples, including mouse brain, human PDAC, and Drosophila embryos, has revealed specific spatial gene expression patterns and provided new insights into cellular organization. Overall, glmSMA represents a powerful and versatile tool for unraveling spatial transcriptomic complexities, offering unprecedented precision in spatial gene expression analysis and making significant contributions to our understanding of tissue architecture.

**Figure 1.**
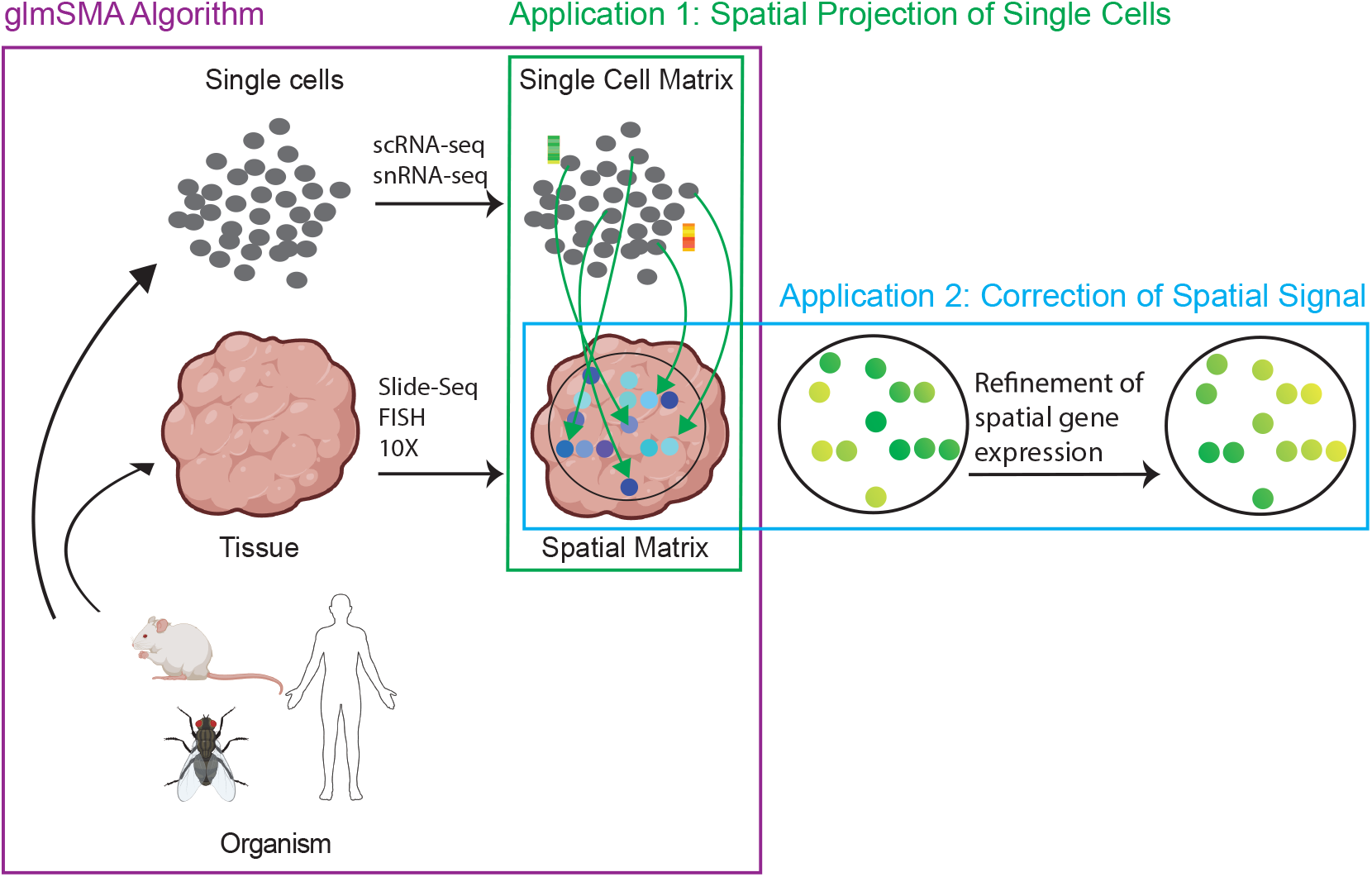
Overview of glmSMA. GlmSMA is designed to estimate the linear relationship between spatial gene expression at specific locations and cellular expression of marker genes, incorporating both L1 and generalized L2 norms for optimal mapping precision. The L1 norm encourages sparsity, ensuring that most single cells are assigned to a small, defined patch within the reference atlas. The generalized L2 norm, meanwhile, promotes smoothness in cell distribution by leveraging anatomical structures and spatial coordinates. This refined mapping can then be used to enhance original spatial patterns.

### GlmSMA successfully map single cells back to mouse cortex based on Spatial-transcriptiomcs data

Spatially resolved gene expression enables researchers to explore how gene expression varies across different regions of a tissue, providing valuable insights into tissue architecture and organization. However, the spatial resolution of 10x Genomics Visium is limited to approximately 55 µm per spot, which is relatively coarse compared to more advanced imaging-based methods like MERFISH or seqFISH^5^. This resolution may not capture finer spatial features, such as single-cell details or subtle gradients in gene expression. As a result, Visium may struggle to distinguish between closely related cell types that are in close proximity within the same tissue region. GlmSMA can address these limitations by incorporating scRNA-seq data. In this approach, individual cells are assigned to small spatial patches based on the 10x Visium dataset, allowing for more precise mapping and better resolution of cellular features within the tissue. To evaluate glmSMA’s performance in accurately mapping cells to their original locations using Spatial Transcriptomics data, we conducted a comparative analysis with CellTrek^12^ and novosparc^10^. Our goal was to map 4,785 single cells, comprising 13 major cell types (Smart-seq2^17^), across 1,075 spots (Visum, 10x Genomics) in the mouse cortex. This dataset spanned six anatomical regions: L1, L2, L3, L4, L5, and L6. We then compared the mapping accuracy of the three algorithms.

GlmSMA accurately mapped 90% of the 4,785 cells back to their original anatomical structures with high confidence (Fig. 2a). For instance, L4 cells were correctly assigned to L4, and L2/3 IT cells were accurately mapped to L2 and L3^31^. Each spot (pixel) contains between 1 and 20 cells. In comparison, CellTrek^12^ mapped approximately 60% of cells with high confidence, but the remaining cells were excluded due to low confidence scores (Supplementary Fig. 1). Novosparc^10^ also mapped 90% of the cells, but its accuracy was lower; for example, around 100 L4 cells were mistakenly assigned to L2 or L3 (Supplementary Fig. 2 and Fig. 2c). Additionally, Novosparc tended to distribute cells more uniformly across the six layers, which does not reflect the actual spatial distribution.

**Figure 2.**
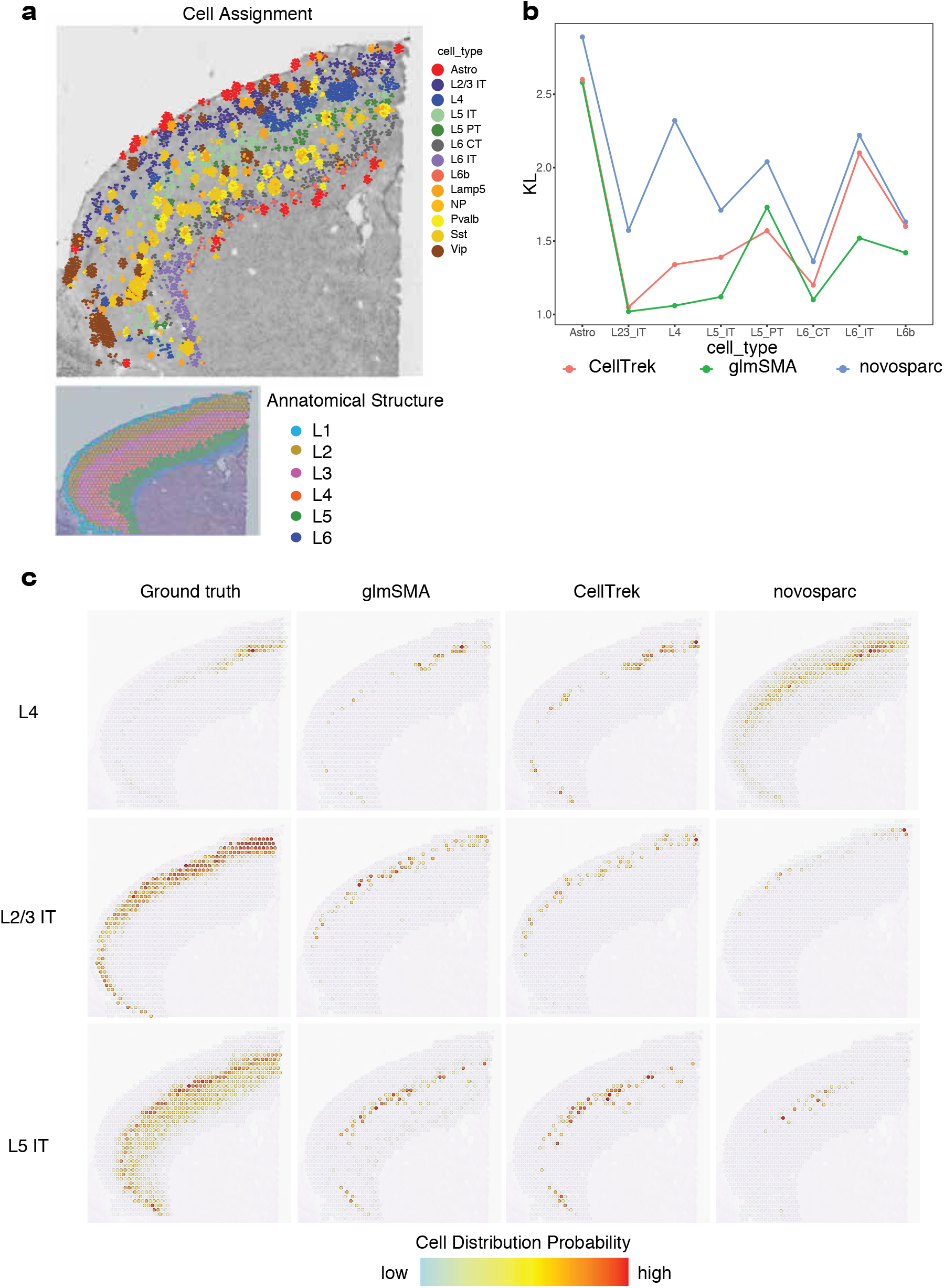
glmSMA reconstruct spatial organization in the mouse cortex. a, Comprehensive Mapping of Cells in the Mouse Brain Cortex: Mapping of 4,785 cells, representing 13 distinct cell types, within the mouse brain cortex, visualizing the spatial organization of these cellular populations. b, KL-Divergence Analysis Across Methods: A comparison of Kullback-Leibler (KL) divergence values between predicted cell type distributions and ground truth distributions across three methods. Results show that glmSMA achieves superior prediction accuracy for most cell types, aligning closely with actual distribution patterns. c, Comparison of predicted and ground truth cell distributions in specific cell types.

To quantify differences among the three algorithms, we calculated the KL-divergence between the predicted and ground truth distributions. Since we lacked exact ground truth for single-cell locations, we approximated the spatial cell distribution based on the spatial transcriptomics data and Allen Brain Atlas^19,20^ using the Seurat package^18^. Subsequently, we evaluated the predicted cell distributions against the reference atlas, which served as an alternative ground truth. Based on the KL-divergence scores, glmSMA outperformed the other two algorithms for most major cell types, though CellTrek slightly outperformed glmSMA for Astro cells (Fig. 2b). Furthermore, the spatial patterns predicted by glmSMA and CellTrek for key cell types, such as L4 and L2/3 IT cells, closely resembled the ground truth distribution (Fig. 2c). In conclusion, glmSMA demonstrates its applicability in spatial transcriptomics data, showing better performance to CellTrek and novosparc, and accurately recapitulating the spatial distribution of individual cells.

### GlmSMA accurately predict cells enrichment in intestinal villus and human PDAC-A sample without spatially informative genes

Spatial-omics techniques, especially transcriptomics, may suffer from low sensitivity, leading to incomplete or sparse data. This occurs because only a subset of genes may be detected in each spatial spot (e.g., in spatial transcriptomics), especially for lowly expressed genes. Incomplete data reduces the ability to accurately characterize gene expression across the entire tissue and may miss critical molecular features or interactions. To investigate the accuracy of glmSMA in tissues with limited spatially informative genes, we applied the method to the human PDAC-A sample^9^. Our objective was to map 1,926 cells, representing 11 cell types, back to 428 locations across four anatomical regions: ductal epithelium, cancer region, stroma, and pancreatic tissue. Evidence suggests that as the physical distance between spots increases, the expression of marker genes becomes more distinct (Fig. 3a). However, Moran’s I score analysis revealed that most of the highly variable genes in this dataset did not exhibit clear spatial patterns. In fact, adjacent spots within a single anatomical region often showed markedly different gene expressions (Fig. 3b and Supplementary Fig. 3). Despite the absence of spatially informative genes, glmSMA successfully mapped the major cell types to their original locations, as confirmed by the ground truth from the study (Fig. 3c). As expected, ductal-like cells were primarily enriched in the ductal epithelium, cancerous cells were predominantly found in the cancer region, and acinar cells were most abundant in the pancreatic tissue. These results align with previous research^9^.

**Figure 3.**
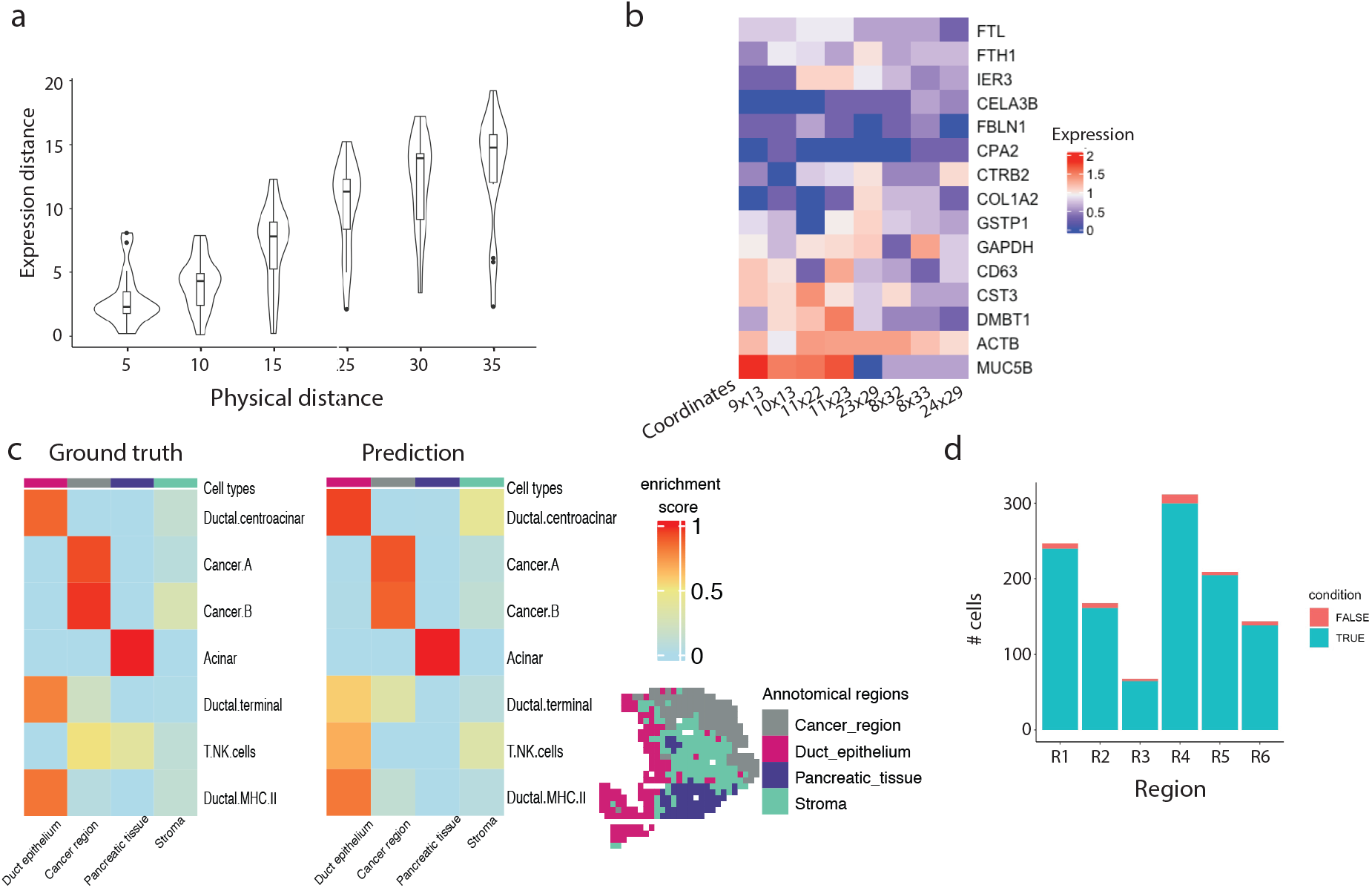
Cell enrichment in the intestinal villus and human PDAC-sample lacking spatially informative genes. a, Violin plots illustrate the pairwise expression differences between spots. Expression differences increase as the physical distance between spots grows. b, Marker gene expression levels across different locations. Adjacent spots can exhibit significantly different expression levels. c, The enrichment of cell types in different anatomical structures. The predicted cell enrichment closely aligned with the ground truth for most cell types. d, The number of cells assigned to each region in intestinal villus.

Next, we investigated another tissue—intestinal villus—with similar challenges of lacking spatially informative genes^21^. Using the top 100 highly variable genes, we were able to correctly map 97% of the cells to their original layers (Fig. 3d). These findings demonstrate that glmSMA can perform well even when relying solely on highly variable genes.

### GlmSMA effectively assigns single cells to their original locations within the 3D surface of the Drosophila embryo

As technologies advance, 3D spatial transcriptomics is anticipated to become increasingly accessible and applicable across various research fields^26^. Our next objective is to assess whether our algorithm can accurately function in 3D space. Since most spatial transcriptomics datasets are derived from 2D tissue slices, we opted to use a simplified 3D surface model for this study. To establish a reference profile, we used an expression atlas of 60 out of 84 marker genes in developmental stage 5, sourced from the Berkeley Drosophila Transcription Network Project (BDTNP)^23,23^. Embryos in the BDTNP database were collected at this developmental stage, with fluorescent staining applied to label nuclei and the expression patterns of marker genes.

We mapped 1,297 single cells onto 3,039 locations in 3D space, with these cells distributed across the entire model (Fig. 4a). Due to the absence of ground truth for single cells, we adopted an alternative strategy: comparing the spatial differential patterns of well-known marker genes with our reconstructed patterns. Reconstructed patterns were generated by assigning scRNA expression data to their respective locations. Our algorithm, glmSMA, effectively captured the spatial expression patterns of marker genes (Supplementary Fig. 4). For instance, the reconstructed ftz patterns displayed segmental stripes similar to the ground truth, while sna expression was high on the ventral side and low on the dorsal side, as expected (Fig. 4c).

**Figure 4.**
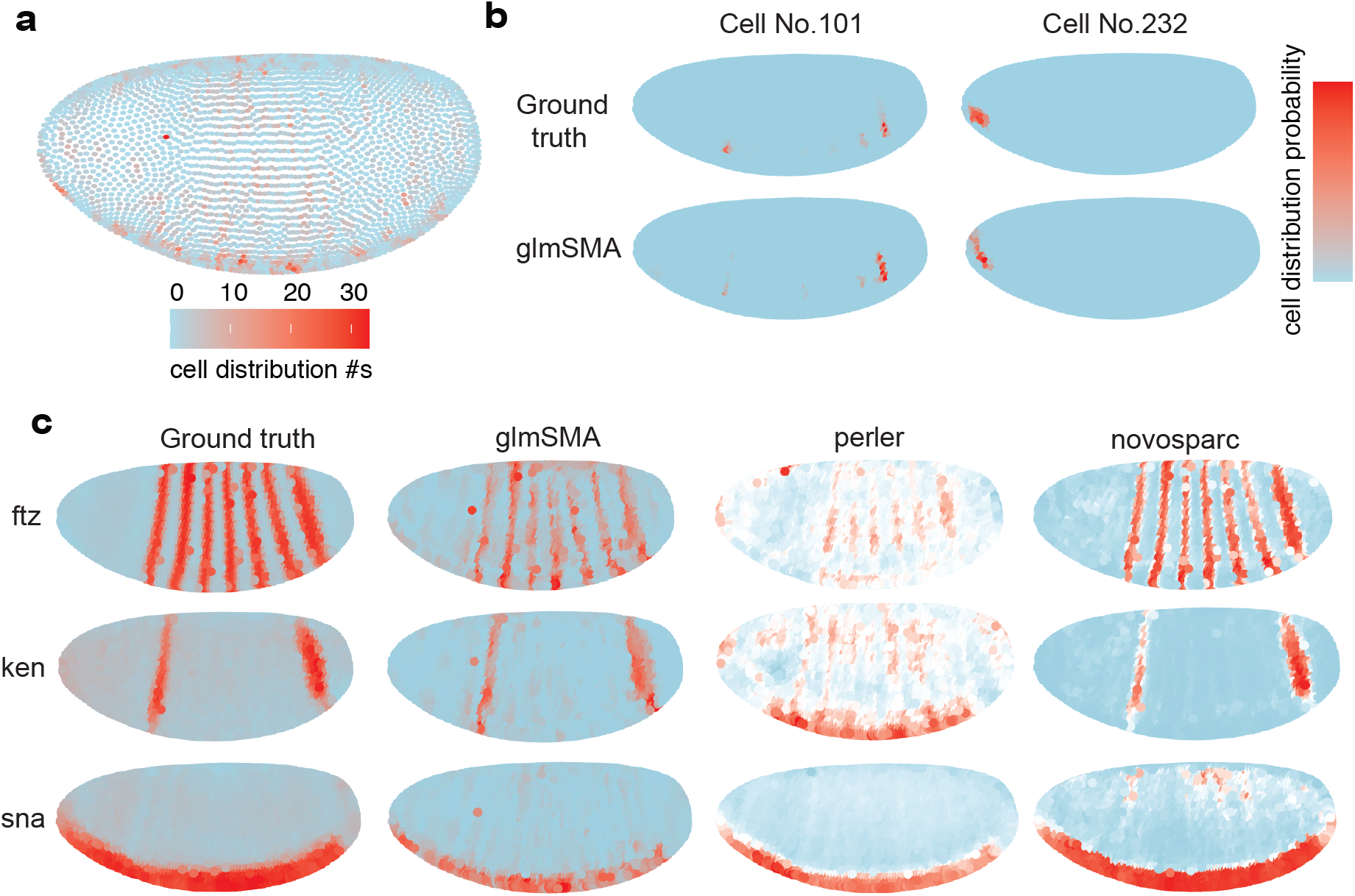
Reconstruction of drosophila embryo. a, Overall 1,297 cell assignment in drosophila embryo. Multiple cells can be assigned to same locations. b, Individual cell assignment. Each cell will be assign to a small patch. c, Comparison of glmSMA, perler and novosparc results for reconstruction of marker genes in a drosophila embryo.

We further compared our algorithm to other methods. Both glmSMA and novosparc produced results consistent with the ground truth, while perler^24^ struggled to replicate the patterns for some marker genes, such as ken (Fig. 4c). We also evaluated cell assignments on an individual level. When comparing our cell assignments to those generated by novosparc, used here as a reference, we found that our assignments were highly consistent (Fig. 4b & Supplementary Fig. 5). For example, cell 232 was accurately assigned to the embryo’s frontal region. By incorporating L1 and L2 regularization, cells were encouraged to form more cohesive patches rather than being scattered across multiple locations (Supplementary Fig. 7). Overall, these results suggest that glmSMA performs well in assigning single cells within a 3D spatial framework.

### GlmSMA successfully maps single cells within the mouse brain at high resolution

Slide-seq is a powerful tool for spatial transcriptomics, offering high-resolution data and large-scale spatial maps of gene expression. However, its capture rate is relatively low, leading to a sparser dataset. To demonstrate glmSMA’s versatility in high-resolution data, we evaluated its performance in mapping cells within the mouse cerebellum. First, we established a reference atlas using Slide-seq, containing a sagittal section with 46,376 spatial locations, each containing 1–1.5 individual cells and covering 19,782 genes^10^. To simulate scRNA-seq data, we coarse-grained the cerebellum by binning neighboring locations in the reference atlas, following a method similar to that used by Nitzan et al. Each resulting patch corresponds to a median spatial diameter of approximately 50 µm, providing a practical estimate of the mapping resolution in the simulated data. After simulation, we tested glmSMA’s ability to map the simulated 7,724 cells back to the original 46,376 locations using the top 100 highly variable genes. As expected, glmSMA accurately assigned 99% of the simulated cells to their original locations (Fig. 5a).

**Figure 5.**
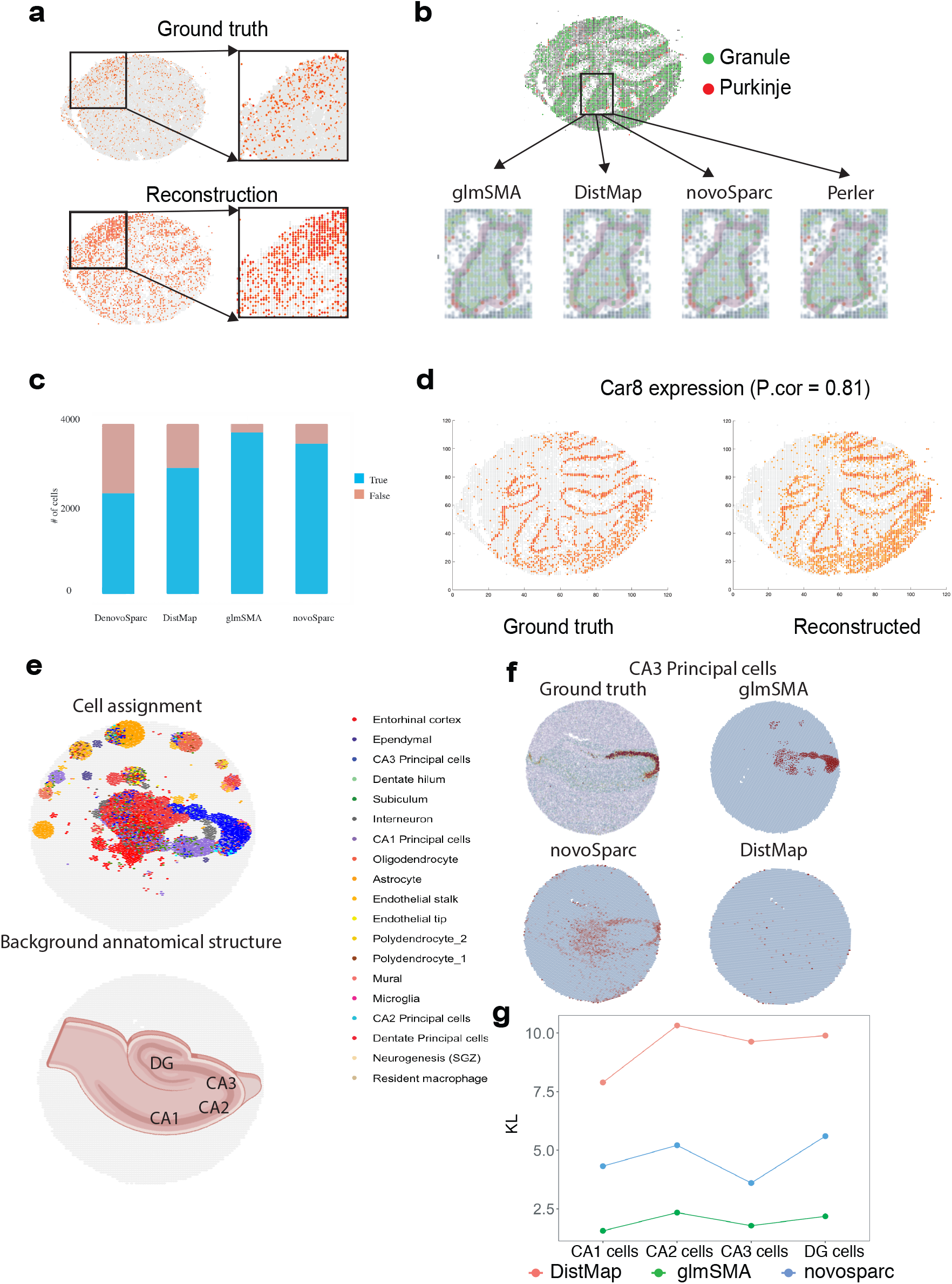
Reconstruction of mouse cerebellum. a, Simulation results of glmSMA for cell assignment in the mouse cerebellum. b, Comparsion of glmSMA, DistMap, novosparc and perler results for Granule cell and Purkinje cell assignments in the mouse cerebellum. c, Comparsion of glmSMA, DistMap denovosparc and novosparc results for Granule cell assignment in the specific anatomical region. d, Reconstruction of Car8 gene using glmSMA with the reconstructed pattern consistent with the ground truth. e, Overall of single cell (19 cell types) assignment in the mouse hippocampus. f, Comparison of predicted and ground truth cell distributions for CA3 Principal cells. g, KL divergence values between predicted cell type distributions and ground truth distributions across three methods. GlmSMA outperformed others.

Next, we assessed glmSMA’s performance against other mapping algorithms. We obtained scRNA-seq data from DropViz^25^, which includes 5,138 cells from the mouse cerebellum, comprising 4,103 granule cells and 310 Purkinje cells. The majority of granule and Purkinje cells were correctly assigned to their anatomical structures, with expression patterns of marker genes, such as Car8, aligning well with the ground truth (Fig. 5b & 5d). When comparing prediction accuracy across algorithms, glmSMA showed superior performance. While other algorithms often misassigned Purkinje cells into granule regions, glmSMA achieved higher accuracy, correctly mapping 3,798 out of 4,103 granule cells to their original anatomical structures (Fig. 5b & 5c). Most Purkinje cells were also accurately assigned, surrounding the granule cells as expected. Additionally, glmSMA demonstrated robust performance on the Slide-seq dataset from the mouse hippocampus (Supplementary Fig. 6). These findings underscore glmSMA’s strength in providing accurate spatial mapping in complex tissues like the cerebellum.

To further assess performance on Slide-seq data with complex cell compositions, we applied glmSMA to the mouse hippocampus. The scRNA-seq dataset contains approximately 50,000 cells, and the Slide-seq dataset includes 7,338 spots after coarse-graining, with each spot representing ∼5–8 cells. The effective spatial resolution of each coarse-grained spot corresponds to a diameter of approximately 50 µm. A total of 19 cell types were successfully mapped back to their original locations. As expected, CA1, CA3, and dentate principal cells were accurately localized to the CA1, CA3, and DG regions, respectively (Fig. 5e). Due to the absence of ground truth for single-cell positions, we estimated cell-type distributions using the reference atlas as an alternative ground truth. We then compared our predicted distributions to this reference and observed strong consistency, particularly for CA3 cells (Fig. 5f). Next, we accurately reconstructed the expression patterns of marker genes based on the mapping results (Supplementary Fig. 8). Finally, we benchmarked glmSMA against NovoSpaRc and DistMap using KL-divergence (Fig. 5g). GlmSMA outperformed both methods across all major cell types, demonstrating its robustness and accuracy in high-resolution spatial transcriptomics data.

## Discussion

Here, we proposed a new method to map the individual cells back to their original locations by combining reference atlas and scRNA-seq data. We take advantage of both techniques in measuring gene expression levels in different tissues. ScRNA-seq can provide comprehensive expression profiles of many individual cells, but location information is missing due to experimental procedure. Slide-seq or FISH-based technology can capture expression profiles at cellular level but only for a limited number of genes. Although the Slide-seq technique can measure the whole genome in the desired tissue simultaneously, only a few numbers of genes are available for the downstream analysis after normalization and filtering.

Existing single-cell assignment methods face notable limitations. For instance, novoSparc primarily emphasizes simulated datasets, as seen in their focus on the simulated dataset. In the mouse cerebellum, rather than utilizing corresponding scRNA-seq data for spatial-omics information, they opted to simulate scRNA-seq data from the Slide-seq dataset. However, novosparc demonstrated suboptimal performance on real datasets, such as the mouse cortex, where the reconstructed spatial expression patterns of marker genes failed to fully reproduce the original patterns evident in ISH images. Another method, DistMap, oversimplifies mapping problems by binarizing expression profiles in both scRNA-seq data and the reference atlas. While effective in relatively simple tissues, the performance of this algorithm significantly diminishes in more complex tissues, such as the mouse brain. On the other hand, Celltrek primarily focuses on spatial transcriptomics data from 10x Visium, requiring a Seurat object as input. In terms of mapping results, it automatically removes at least 30% of cells with low confidence in most cases. In contrast, our method maximizes the relationship between cell expressions and locations. With enough number of marker genes, we can accurately place most cells into correct small patches. While achieving pinpoint accuracy for each individual cell in certain complex tissues may pose challenges, our approach consistently and accurately assigns them to relatively small regions.

Other established methods for mapping single-cell data into specific locations are limited to low resolution. For example, MIA and HMRF^27^ are two methods focusing on locating cell types into regions. Instead of mapping thousands of cells back to several positions, they only aimed to map tens of cell types into large domains. Information on individual cells were lost after the mapping because they merged the expression profiles in the same cell type. In addition, their methods cannot be applied in locating cells back to small patches, but our algorithm can be used to locate cells into domains, demonstrated in the mouse cortex dataset. In the mouse cortex, we successfully mapped cells back to 6 large domains with high accuracy. Therefore, our algorithm provided accurate and high-resolution mapping for the scRNA-seq data compared with the existing low-resolution methods.

Our method to map individual cells back to their original locations also has limitations. First, the performance of our algorithm highly relied on the selection of marker gene sets. Due to the cost of building a reference atlas using FISH-like technology, we may not obtain enough number of gene candidates. However, this can be solved by preprocessing the scRNA-seq or bulk-RNA seq data^28^. The preprocessing step can help to shrink the size of marker gene candidates. Second, the current technology of generating reference atlas, such as Slide-Seq, remained significant defects. The data quality needed to be better improved because expression profiles of lots of well-known marker genes cannot be captured^29^. Despite limitations, our method still achieved good performance in integrating the scRNA-seq data and reference atlas. Most of the individual cells can be correctly mapped to several unique locations.

Most current methods, including glmSMA, perform better on major cell types. This is largely because major cell types typically have well-defined marker genes. In contrast, rare cell types often lack distinct markers and are represented by only a small number of cells, making accurate identification and mapping more challenging.

Combined with the strength of both scRNA-seq and Spatial-omics techniques, our analysis and reconstruction of spatial expression patterns identified specific cell organizations in physical space under the assumption that cells with closer physical distance are more likely to have more similar expression profiles. Reconstructed spatial patterns of specific marker genes in the mouse brain have been confirmed by the ISH images from Allen Brain Atlas and Hipposeq^30^. Also, our mapped results provided an alternative way to impute the low-quality data in Slide-seq.

## Conclusions

GlmSMA is a groundbreaking linear regression model meticulously crafted to seamlessly fuse scRNA-seq reference data with spatial transcriptomics (ST) data obtained from various experimental platforms. Through a comprehensive benchmarking process encompassing simulated and in situ datasets, glmSMA demonstrates unparalleled performance, successfully assigning single cells to their precise original locations. The application of glmSMA in mapping single cells back to their origins has revealed novel insights, underscoring its effectiveness as a versatile and potent tool for unraveling the intricacies of spatial transcriptomics.

## Code availability

Code related to this manuscript can be found at https://github.com/liuzdlab/glmSMA/.

## Acknowledgement

This study was supported in part by the Cancer Prevention and Research Institute of Texas (CPRIT) award RP240131 (to C.G., H.C., and Z.L.).

## Author Contributions

Chaohao and Zhandong designed the glmSMA; Chaohao implemented the glmSMA, and carried out all the analyses, under Hu’s co-supervision. Chaohao and Zhandong wrote the manuscript with inputs from Hu.

## Author Notes

Competing interests: No competing interests declared.

