## Supplementary figures and images for "A network regularized linear model to infer spatial expression pattern for single cell" (from another version of this preprint: version 2, 2025-07-09)

### Supplemental Figure 1

Supplementary Figure 1: Mouse cortex reconstructed by CellTrek.

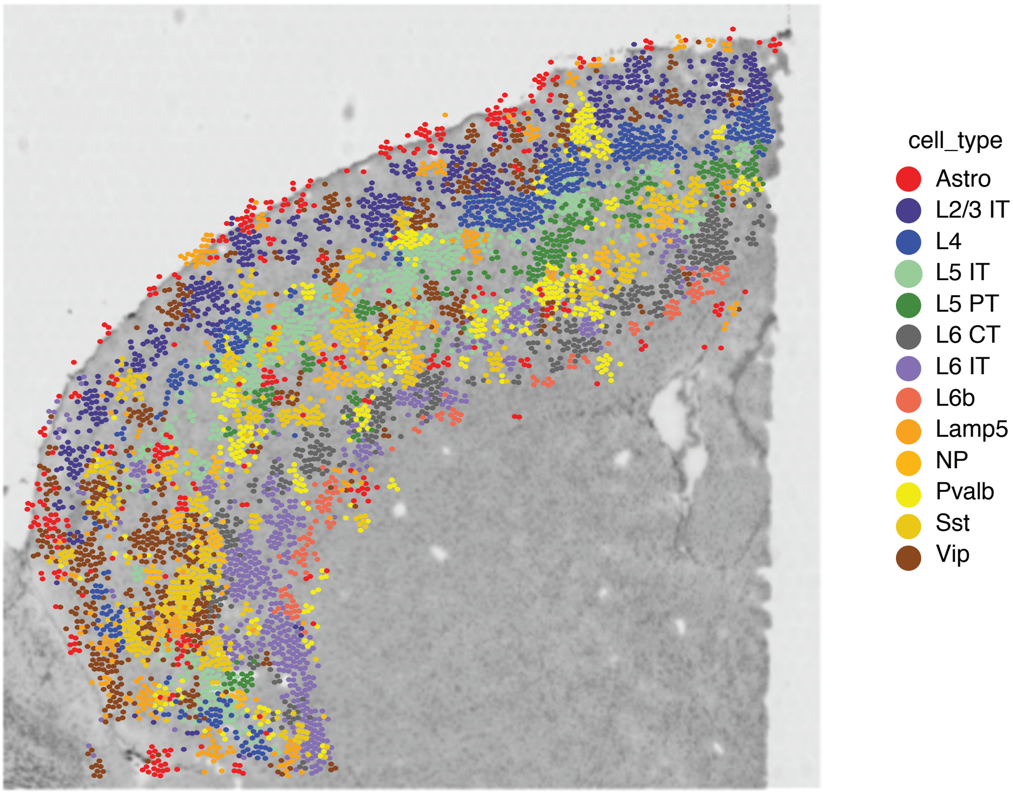


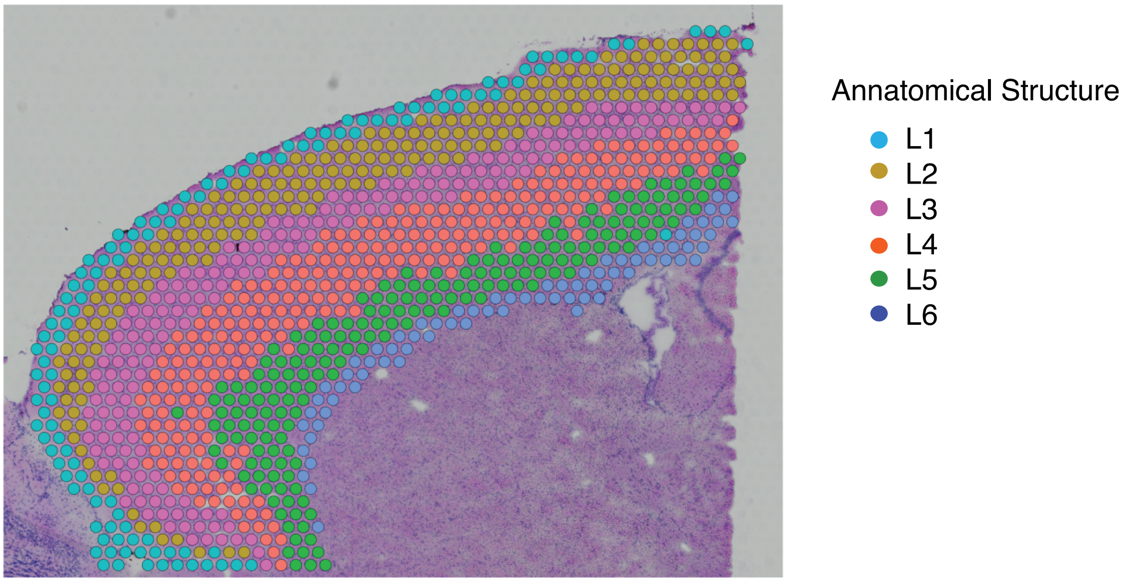

### Supplemental Figure 2

#### Supplementary Figure 2: Mouse cortex reconstructed by novosparc.

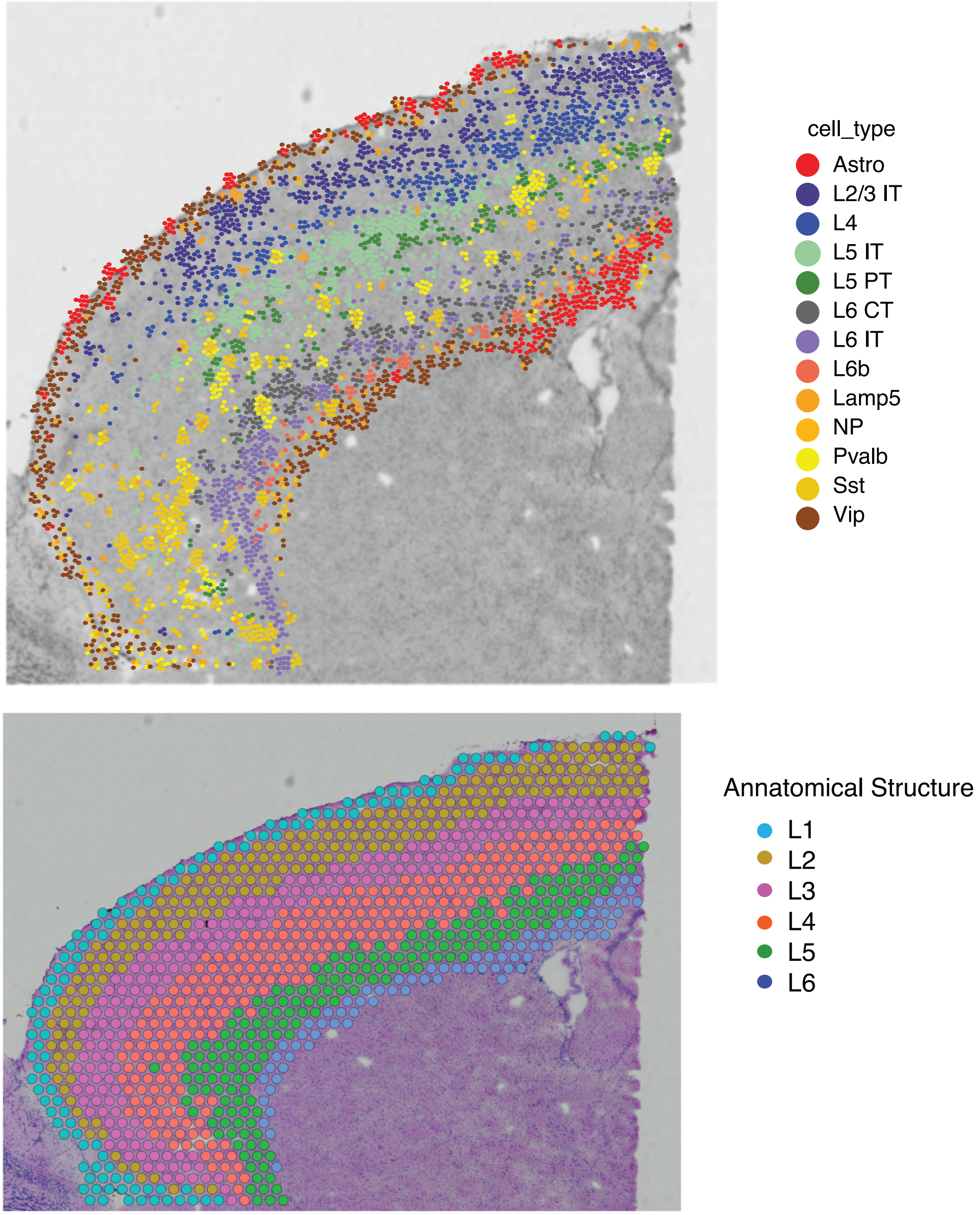

### Supplemental Figure 3

Supplementary Figure 3: Moran's I for markers.

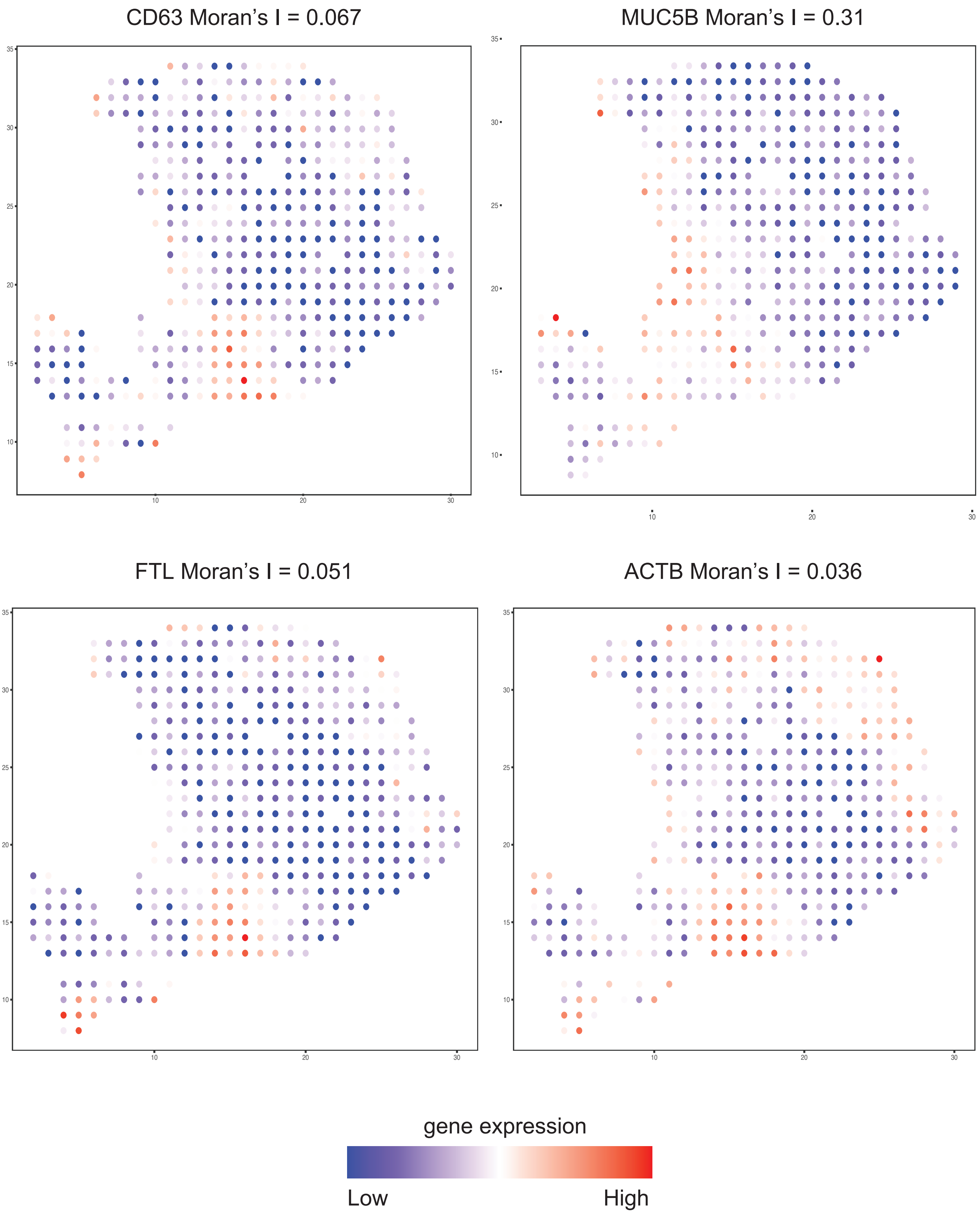

### Supplemental Figure 4

Supplementary Figure 4: Marker genens expression reconstructed by glmSMA.

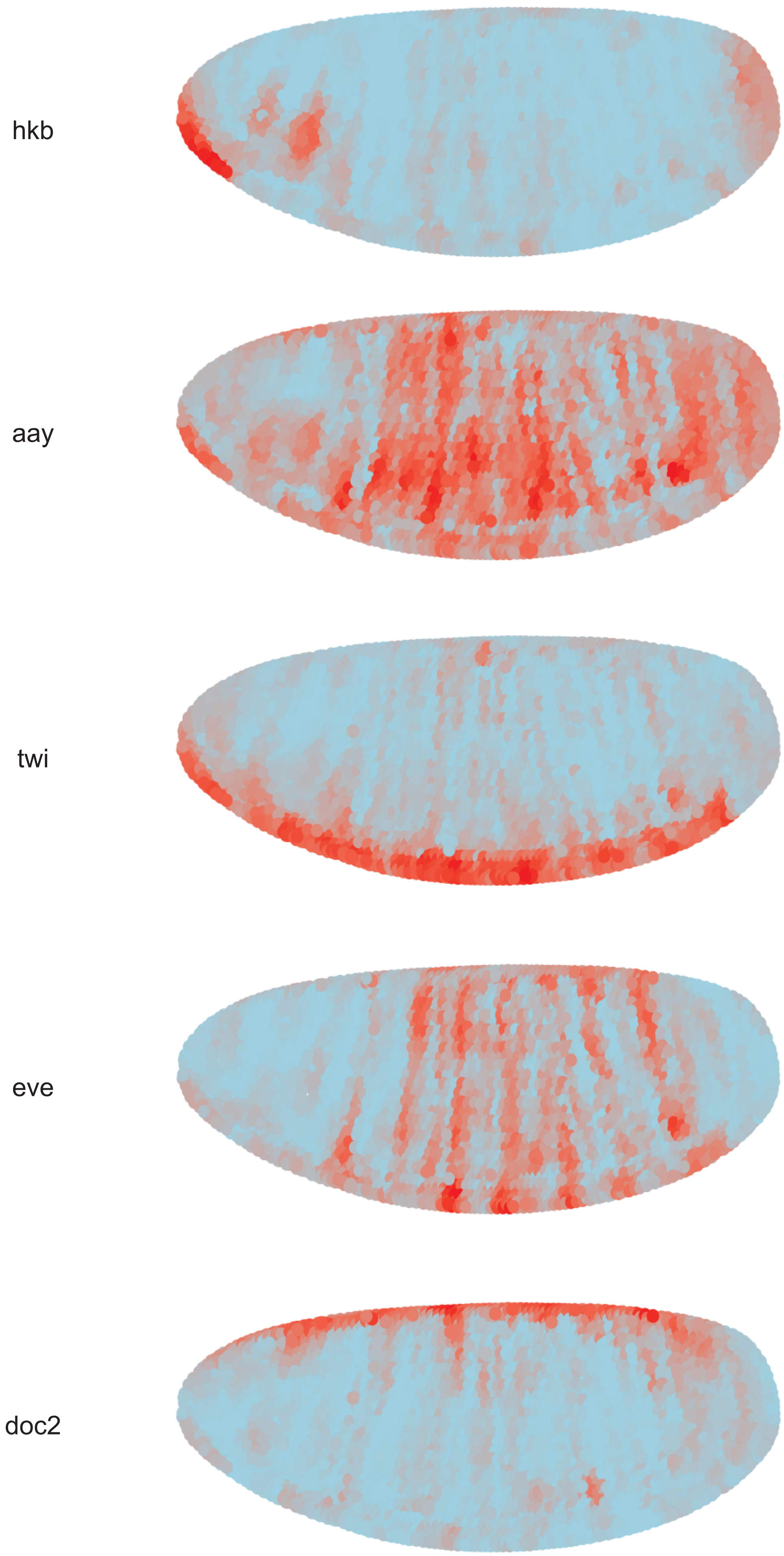

### Supplemental Figure 5

Supplementary Figure 5: Cell Assinment by glmSMA.

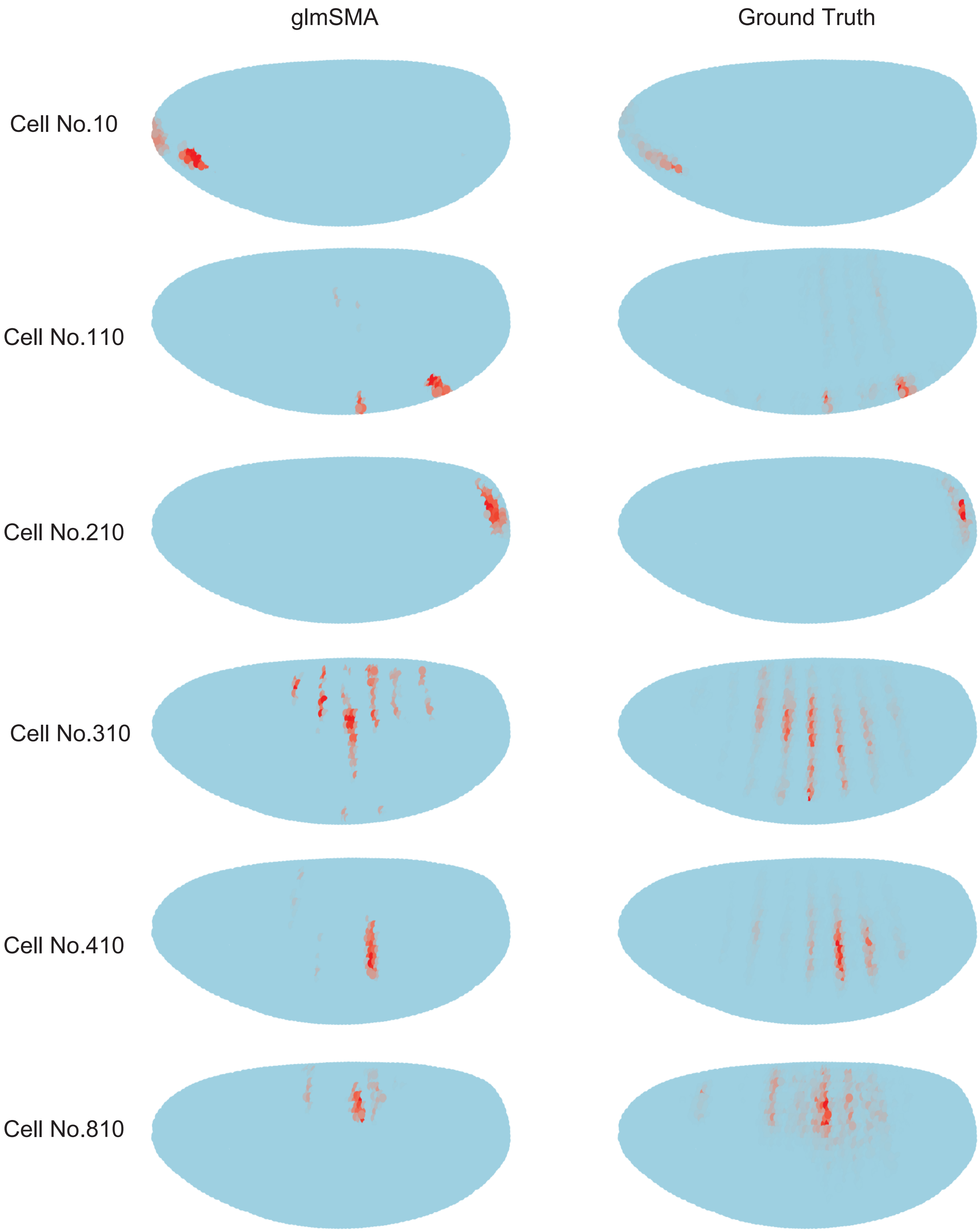

### Supplemental Figure 6

Supplementary Figure 6: Cell assignment in the mouse hippocampus.

a

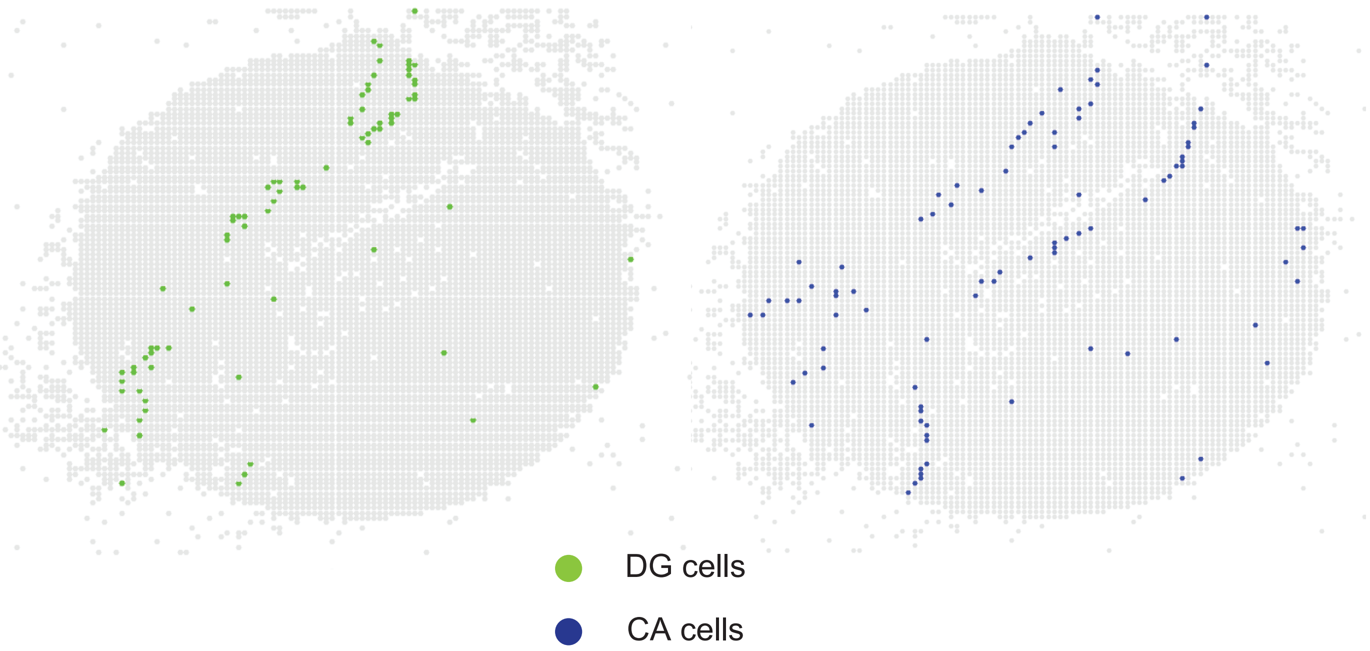


b

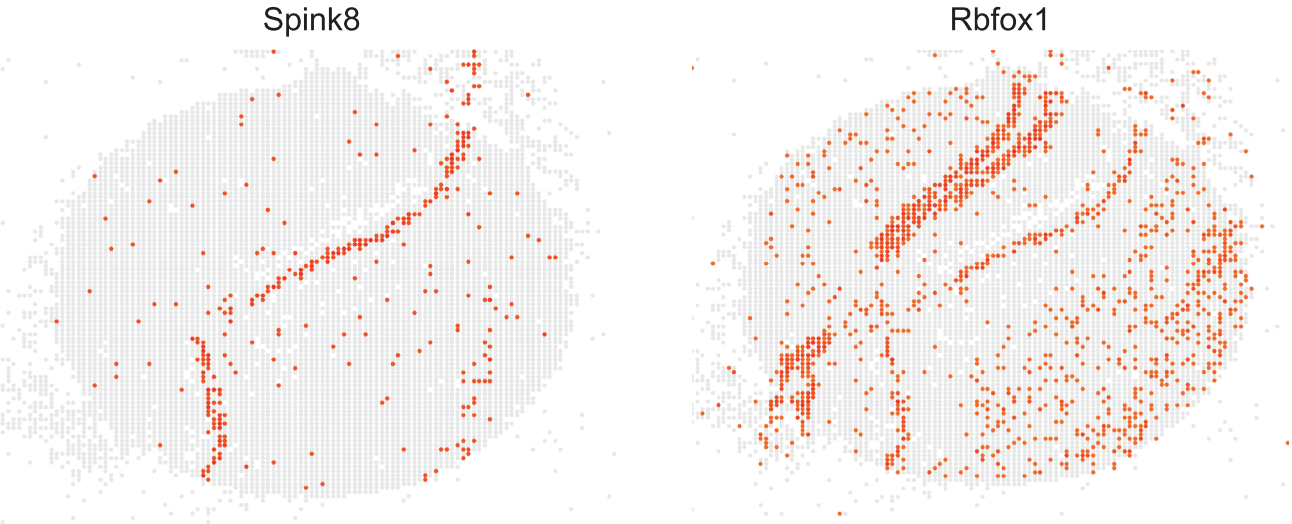

### Supplemental Figure 7

Supplementary Figure 7: Regularization path for cell assignment.

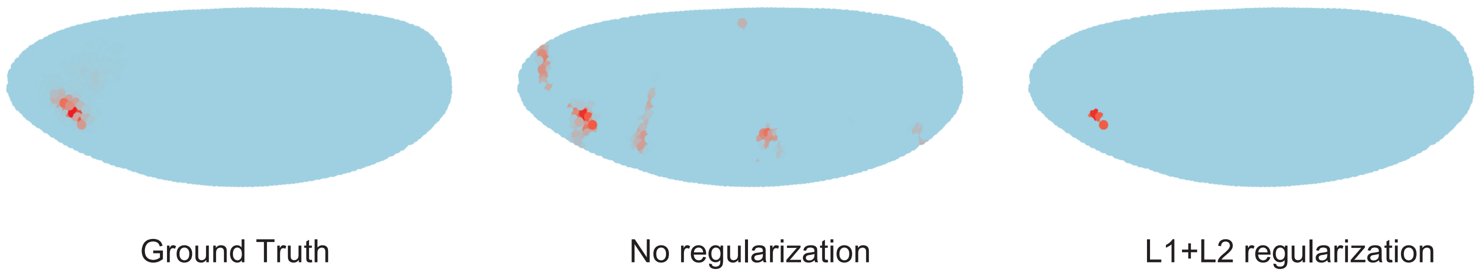

### Supplemental Figure 8

Supplementary Figure 8: Reconstructed marker genes of mouse hippocampus.

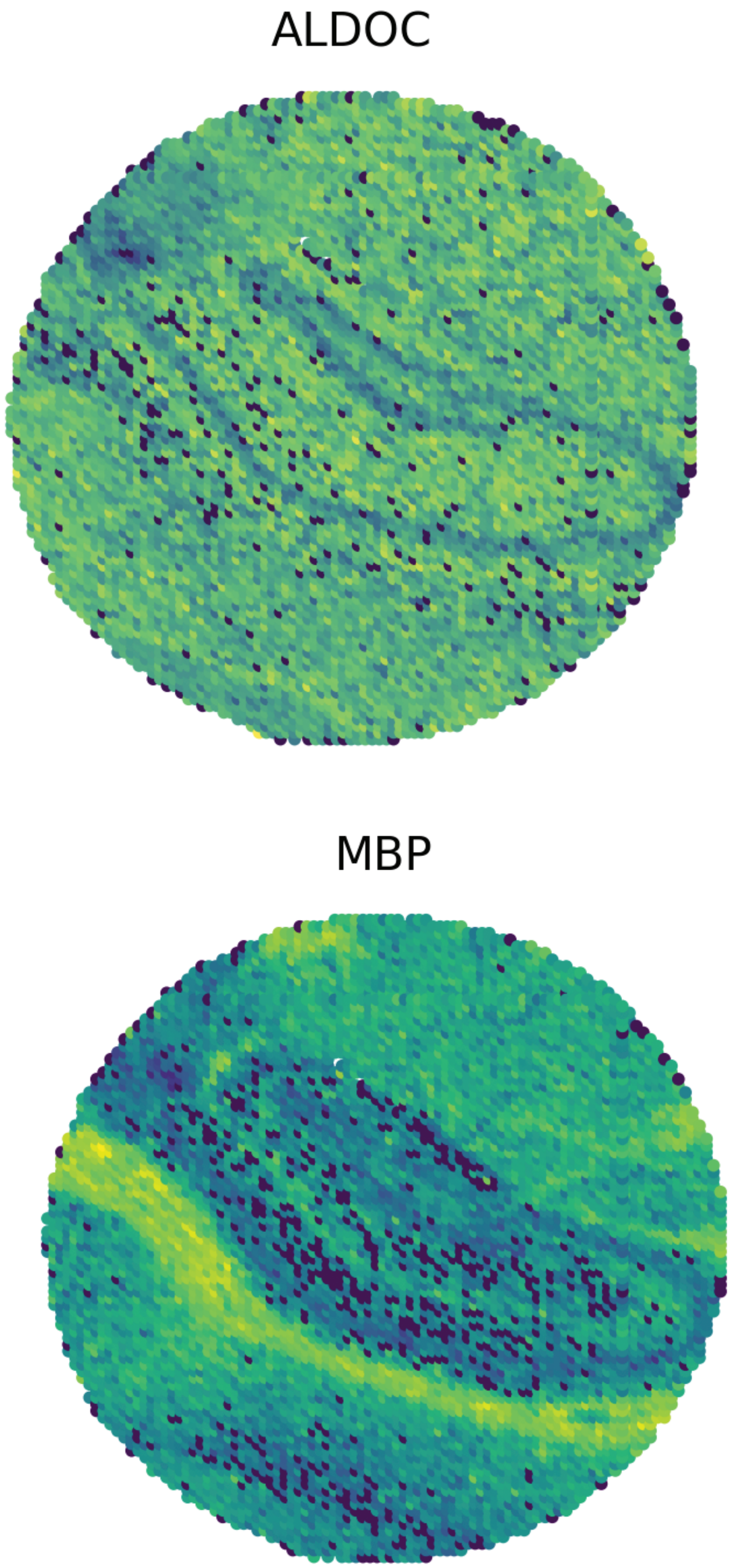

### Supplemental Figure 9

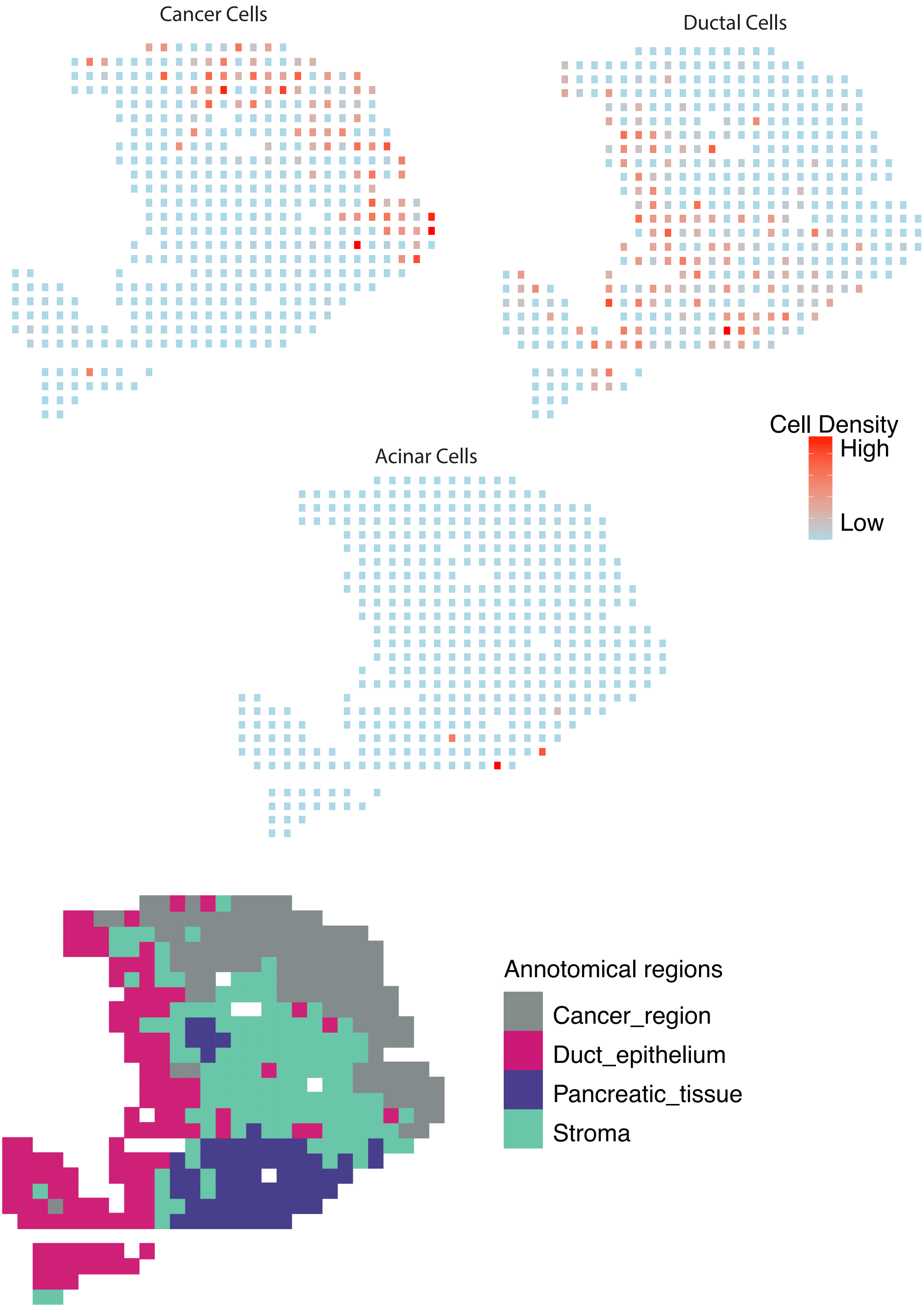

### Supplemental Figure 10

Supplementary Figure 10: Impact of marker gene number on cell assignment.

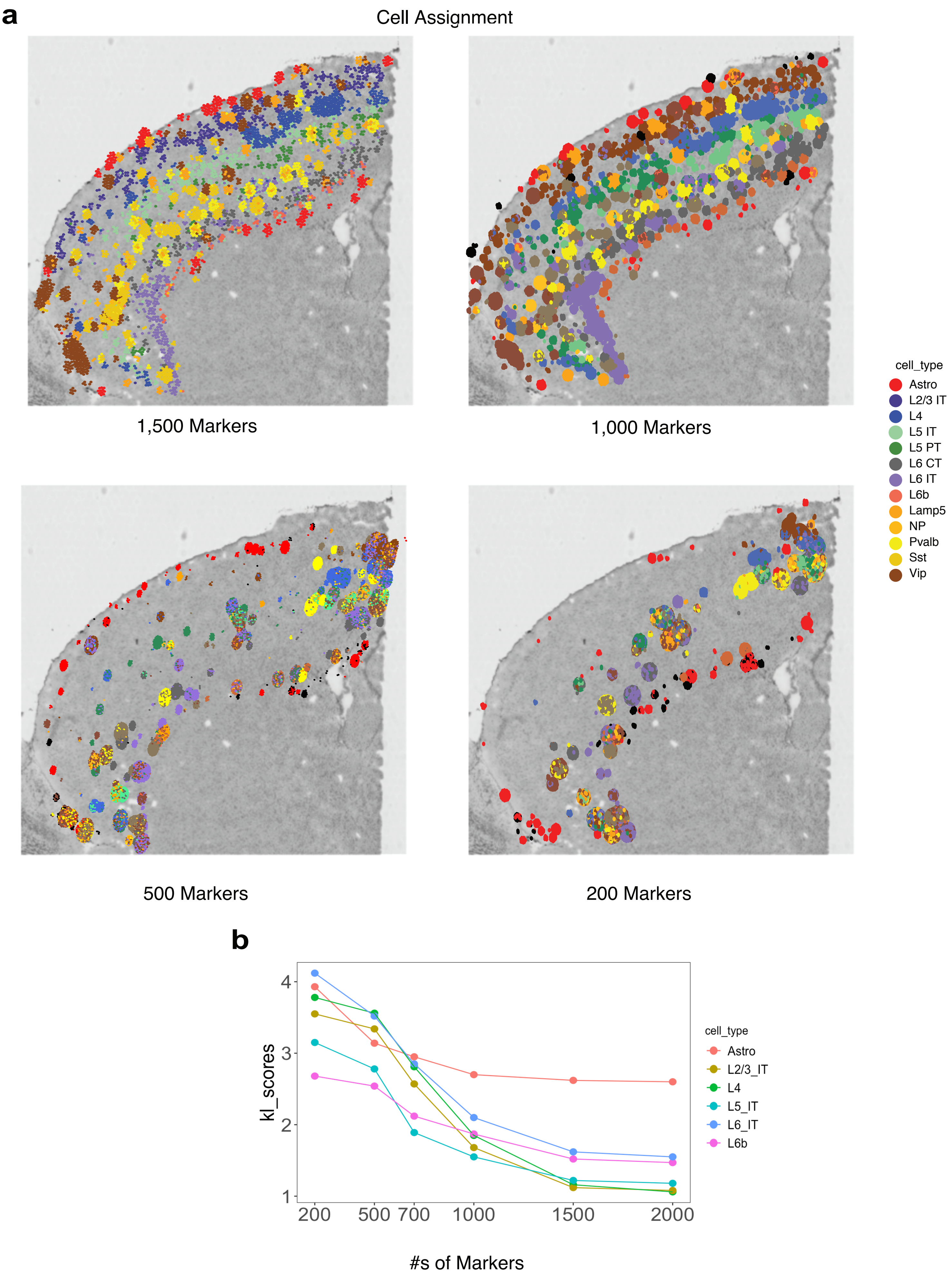
